# The neural basis of cost-benefit trade-offs in effort investment: a quantitative activation likelihood estimation meta-analysis

**DOI:** 10.1101/2022.10.28.513278

**Authors:** Nathaniel Edgar, Kevin da Silva Castanheira, Gary R. Turner, R. Nathan Spreng, Eliana Vassena, A. Ross Otto

## Abstract

Influential theories of cognitive effort-based decision-making suggest that a cost-benefit trade-off guides mental effort allocation and that this trade-off may be reflected in shared neural activity in circuits tracking potential reward and task demand, supporting the idea that. While the dorsal medial prefrontal cortex (mPFC) - and particularly the anterior cingulate cortex (dACC) - has been proposed as a candidate region implementing this computation, it remains unclear whether mPFC/dACC activity tracks rewards and task demand independently or integrates them to reflect effort intensity. Recent accounts posit that the dACC plays a key role in mediating cost-benefit trade-offs. However, empirical evidence remains inconsistent. We conducted a systematic meta-analysis of neuroimaging studies, using the activation-likelihood estimation method to quantify brain activity across 45 studies (*N* = 1273 participants) investigating choices and task performance in reward-guided cognitive control. We observed significant recruitment of the mPFC/dACC, putamen, and anterior insula for processing larger rewards and higher task demands. The mPFC/dACC clusters sensitive to task demands and rewards were anatomically distinct: caudal mPFC/dACC activity tracked increasing task demands, while rostral mPFC/dACC activity tracked increasing reward. Interestingly, caudal mPFC/dACC activity tracked the integration of reward and task-demand, compatible with cost-benefit trade-off theories of dACC function. These findings provide evidence for distinct signals for mental demand and reward in the mPFC/dACC, which are integrated to support the decision to invest mental effort.

## Introduction

How do we decide whether pursuing a reward is worth the mental effort required to obtain it? On one hand, the experience of cognitive effort exertion is aversive (and often avoided), yet, on the other hand, individuals must often engage in effortful thinking to obtain rewards. Consequently, our decisions to engage in cognitively costly processing often present a conflict between two opposing goals: maximizing rewards and minimizing the associated effort costs. To this point, prominent theories of motivated control posit that deciding to allocate cognitive effort requires the integration of the benefits (e.g., rewards) tied to effort exertion, the costs of effort, and the likelihood of successful performance (Frömer et al., 2021; Kurzban et al., 2013; Shenhav et al., 2017; Silvetti et al., 2018). Indeed, a large and growing body of empirical work suggests that our decisions to allocate effort result from an integration of costs and benefits. For example, reward incentives motivate cognitive effort investment (Otto & Vassena, 2021; Westbrook & Braver, 2015), particularly for individuals with low executive function capacity (da Silva Castanheira et al., 2021; Sandra & Otto, 2018). Further, during decision-making, the subjective value of rewards is typically discounted (i.e. decreased) by the effort required to earn these rewards (Chong et al., 2017; Otto & Vassena, 2021). People will even opt for a physically painful sensation over the prospect of exerting high levels of cognitive effort (Vogel et al., 2020).

Influential theories suggests that the medial prefrontal cortex (mPFC) - and particularly the dorsal anterior cingulate cortex (dACC) - plays a role in resolving this cost-benefit trade-off by integrating specific neural signals representing both effort costs and anticipated rewards (Shenhav et al., 2013; Silvetti et al., 2018). The functional role of the dACC has been highly debated ((Ebitz & Hayden, 2016; Shenhav et al., 2013; Vassena et al., 2017, 2020), and the various anatomical labels used to indicated several portions of the mPFC have often hindered clear communication of findings across studies and fields. Despite the wealth of behavioral findings supporting the idea of a cost-benefit tradeoff in mental effort exertion decisions (Otto et al., 2025; Silvestrini et al., 2022), the neural evidence supporting the functional role of mPFC/dACC in coding reward, mental effort costs, and/or the integration thereof remains unclear.

For anatomical clarity—and in light of the broad number of studies where these regions are observed to be co-active (Lopez-Gamundi et al., 2021; Parro et al., 2018)—here we employ the term mPFC/dACC to indicate the portion of cortex that expands along the medial wall, covering the rostral, caudal and dorsal portion of anterior cingulate cortex and extends more dorsally towards the pre-supplementary motor area (pre-SMA). The reason for this definition is the wide amount of studies where these regions are found co-active . In line with the predictions of cost-benefit accounts of dACC function (Shenhav et al., 2013; Silvetti et al., 2018), some studies suggest that the mPFC/dACC encodes both reward prospects and task demands. The anticipation of rewards has been consistently associated with greater activity in the mPFC/dACC, anterior insula, thalamus, and ventral striatum (e.g., the nucleus accumbens and putamen; Bartra et al., 2013; Diekhof et al., 2012; Knutson & Greer, 2008). However, these observed patterns of neural activity were not specific to performance-contingent rewards, suggesting a general role for the mPFC/dACC in encoding reward information. More recently, a meta-analysis by Parro and colleagues (2018) investigated activation patterns underlying performance-contingent reward incentives, finding right-lateralized BOLD activity in the mid-dorsolateral PFC, mPFC/dACC, anterior insula, mid-intraparietal sulcus, and anterior inferior parietal lobule in response to rewards. While both the mPFC/dACC and anterior insula have been linked to subjective (self-reported) motivation on cognitive tasks, only the mPFC/dACC has been found to encode integrated incentive values when performing effortful tasks (Yee et al., 2021).

A parallel line of work has identified regions that encode costs associated with increasing task demands either during preparation for tasks or during task performance. When exerting control, increasing task demands engage the mPFC/dACC, posterior parietal cortex, anterior insula, and prefrontal cortex (Laird et al., 2005; Niendam et al., 2012). When anticipating effortful tasks, activity in the mPFC/dACC has been demonstrated to increases as a function of the subjective value of the chosen course of action (Chong et al., 2017). Similarly, a study jointly investigating rewards and task demands observed increases in mPFC/dACC BOLD activity for both larger rewards and task demands (Vassena et al. 2014). However, this study may be limited in its ability to capture effort discounting processes, as it contrasted two demand levels for which high overall accuracy (>90%). Thus, activity in the dACC could either encode the effort level to be invested—that is, the integrated costs and benefits of effort exertion (Chong et al., 2017; Shenhav et al., 2016; Silvetti et al., 2018)—or simply encode a representation of task demands (Lopez-Gamundi et al., 2021).

Beyond simply tracking rewards and demand, the dACC has also been observed to play a role in learning and monitoring task progress. For example, studies have observed that dACC activity tracks negative consequences of errors like negative feedback (i.e., response errors; Carter et al., 1998; Cole et al., 2009; Ito et al., 2003; Ridderinkhof et al., 2004), and pain (Jahn et al., 2016; Shackman et al., 2011). At the same time, the dACC has plays a role in monitoring the need for cognitive control (Botvinick, 2007; Venkatraman & Huettel, 2012), tracking prediction errors (Alexander & Brown, 2015; Brown & Alexander, 2017; Silvetti et al., 2011), and coordinating effortful control over extended action sequences (Botvinick et al., 2001; Holroyd & McClure, 2015). Together, these findings suggest a role for the dACC in the learning of control signal specifications to obtain rewards or avoid punishment (Shenhav et al., 2016). Converging neurocomputational work on adaptive decision-making proposes that the dACC integrates costs and benefits through a meta-learning mechanism via interactions with catecholaminergic input from subcortical systems (the Reinforcement Meta Learner model; Silvetti et al. 2018). In this computational account, the dACC also contributes to the learning of optimal effort allocation over time (Verguts et al., 2015) and to other adaptive learning dynamics (i.e., control of learning rate, higher-order reinforcement learning, Silvetti et al. 2018). Importantly, most of these perspectives rely on the assumption that, to some extent, the dACC receives input signals indexing reward and cost to compute an integrated quantity (net value) that guides effort decisions. This reward signal is supplied via midbrain dopaminergic input, as extensive work in animals has shown (Haber et al., 2006; Haber & Knutson, 2010). On the contrary, the source and neural representation of the cost signal remain highly debated (Holroyd, 2015; Kurzban et al., 2013; Musslick & Cohen, 2021; Wiehler et al., 2022). Whether the cost of cognitive effort is encoded by dACC, and to what extent a reliable signal representing the control signal intensity based on the integration of the costs and benefits is traceable in dACC activity, remains unclear.

With an extensive landscape of mixed findings, drawing reliable conclusions on the neural representation of rewards and effort remains challenging, likely due to the heterogeneity of task designs across studies (e.g., variations in demand type and reward prospects used). in both the. This variability across studies is especially relevant in the case of inconsistent findings, often leading to extensive debates in the literature—for example, in the case of the dACC (Ebitz & Hayden, 2016; Vassena et al., 2017). Meta-analytic synthesis offers the opportunity to isolate reliable effects of interest, allowing for joint investigation of parametric manipulations of demand and reward levels across multiple studies (Yarkoni et al., 2010). Here, we used the activation-likelihood estimation meta-analytic technique (Eickhoff et al., 2012), synthesizing brain activity across 45 studies, to identify circuits selective to task demand, reward prospect, and the integration of both. This allowed testing whether the predictions of the cost-benefit account of dACC function are supported across diverse manipulations of cognitive effort and reward. By looking across studies that manipulate mental demand level and/or performance-contingent rewards, we could probe whether mPFC/dACC activity is associated with effort investment reflecting an integrated representation of costs and benefits, versus a representation of only costs (or only benefits).

Importantly, these different possibilities lead to contrasting hypotheses regarding neural coding of task demand and reward level in mPFC/dACC (see Figure 1). If mPFC/dACC activity only reflects performance-contingent rewards, BOLD responses should increase monotonically with larger rewards but not higher task demands (see Figure 1, left panel). If mPFC/dACC activity reflects only mental demand level (i.e., effort costs), BOLD responses should increase monotonically with higher task demand levels, but not larger rewards (see Figure 1, middle panel). However, if mPFC/dACC activity reflects the integration of demand and reward that leads to effort investment, BOLD responses should scale with both costs and benefits. For example, in the Expected Value of Control (EVC) model (Shenhav et al., 2013), dACC activity is posited to increase with larger rewards and higher task demands (i.e., scales positively with net value; see Figure 1, right panel; Silvestrini et al., 2022). Little work has jointly assessed the neural representations underlying processing both prospective rewards and cognitive demand overlap.

**Figure 1.**
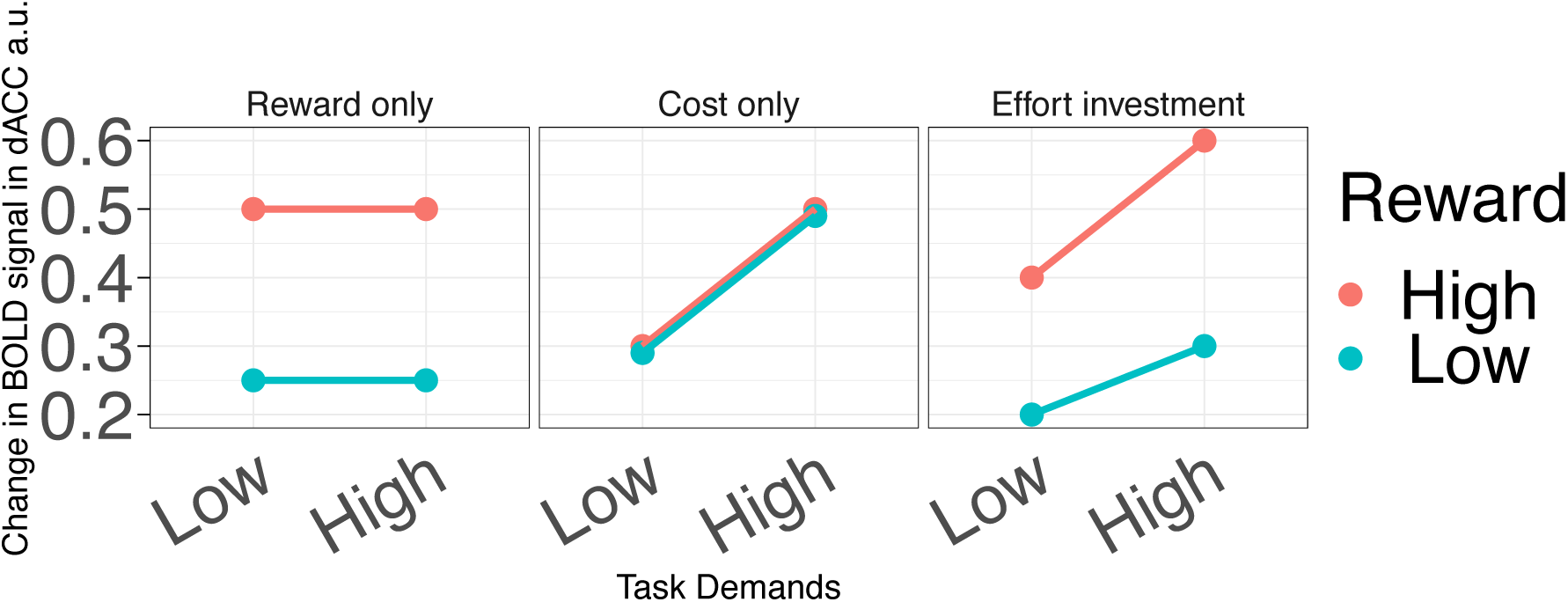
Hypothesized patterns of dACC BOLD signal based on what this signal is thought to reflect: costs only, benefits only, or the effort investment based on the expected value of control. For the reward model, where dACC activity is posited to only reflect reward prospects, BOLD signal is predicted to increase simply as a function of performance-contingent rewards. For the cost model, where dACC activity is posited to only reflect costs, BOLD signal is predicted to increase only as a function of task demands. And for the integration models, where dACC activity reflects the intensity of the control signal or effort to be invested, BOLD signal is predicted to increase in response to higher rewards and higher levels of task demands.

To the best of our knowledge, two previous meta-analyses have examined how brain regions process effort-related costs (i.e., task demands) and integrate them with rewards, finding inconsistent results in partially overlapping regions of the mPFC/dACC. Parro et al. (2018) observed higher mPFC/dACC activity for larger rewards (Parro et al., 2018). In contrast, Lopez-Gamundi et al. (2021) observed increased mPFC/dACC activity for higher demand and reduced activity for higher net value. Interestingly, Lopez-Gamundi also reported an adjacent cluster showing higher activity for higher net-value, raising the question of whether mPFC/dACC reflects task demands, rewards, or an integration of the two.

Additionally, the majority of the studies included in the meta-analyses by Lopez-Gamundi et al. focused on physical effort: whether the observed involvement of mPFC/dACC would extend to mental effort, remains to be clarify. Past behavioral (Lopez-Gamundi & Wardle, 2018), neuroimaging (Chong et al., 2017), and electrophysiological studies in humans (Jiang & Zheng, 2023) as well as pharmacological work in animal models (Hosking et al., 2015) suggests that mental and physical effort may rely, to some extent, on distinct processes (and neural systems). Thus, it is important to clarify whether mental demand and rewards are represented separately, and to what extended they are integrated in a signal compatible with the theoretically hypothesized cost-benefit trade-off, particularly in regions like the mPFC/ACC.

Accordingly, the current meta-analysis aims to disentangle the common and unique patterns of activation observed across several, diverse fMRI studies which independently manipulate reward prospects tied to mental effort exertion, and mental demands across a variety of operationalizations of cognitive demand and reward. Using this approach, we can assess 1) the regions uniquely involved in processing rewards, 2) the regions uniquely involved in processing mental demands and 3) the regions involved in both processes. We further assess the regions which track the interaction between reward and demand signals, encoding the integrated value of mental effort investment. Based on cost-benefit accounts of mental effort-related decision-making (Shenhav et al., 2013; Silvetti et al., 2018), we reasoned that the mPFC/dACC, beyond independently tracking the costs and benefits of mental effort, would integrate these costs and benefits, reflecting the effort level deemed worthy of investing.

## Materials and Methods

### Literature Search

We conducted a systematic review of functional magnetic resonance imaging (fMRI) cognitive control studies which experimentally manipulated either available rewards, (mental) task demand, or both. Our literature search and exclusion process are depicted in the flow chart in Figure 2. We searched for articles published prior to February 3^rd^, 2025, on the online databases PubMed/MEDLINE, Web of Science, and PsychINFO, with an “all fields” search matching the following search string: ("REWARD*" OR "MONETARY INCENTIVE*" OR "MOTIVAT*" OR "INCENTIV*") AND ("COGNITIVE EFFORT" OR "MENTAL EFFORT" OR "COGNITIVE CONTROL" OR "EXECUTIVE FUNCT*" OR "WORKING MEMORY" OR "INHIBIT*" OR "SET SHIFTING" OR "SET-SHIFTING" OR "TASK SWITCHING" OR "TASK-SWITCHING" OR "LOAD" OR ”COGNITIVE LOAD" OR "DIFFICULT*" OR "EFFORT* " OR " DEMAND* ") AND ("FMRI" OR "FUNCTIONAL MAGNETIC RESONANCE IMAGING" OR "BRAIN IMAGING" OR “MRI”) AND ("HUMAN*" OR "PARTICIPANT*" OR "ADULT*” OR “SUBJECT*”). This search yielded 3849 articles. We further included 82 articles which were obtained from manually searching the reference list of previous coordinate-based meta-analyses on either reward processing or effortful control (see Figure 2) (Diekhof et al., 2012; Laird et al., 2005; Lopez-Gamundi et al., 2021; Parro et al., 2018).

**Figure 2.**
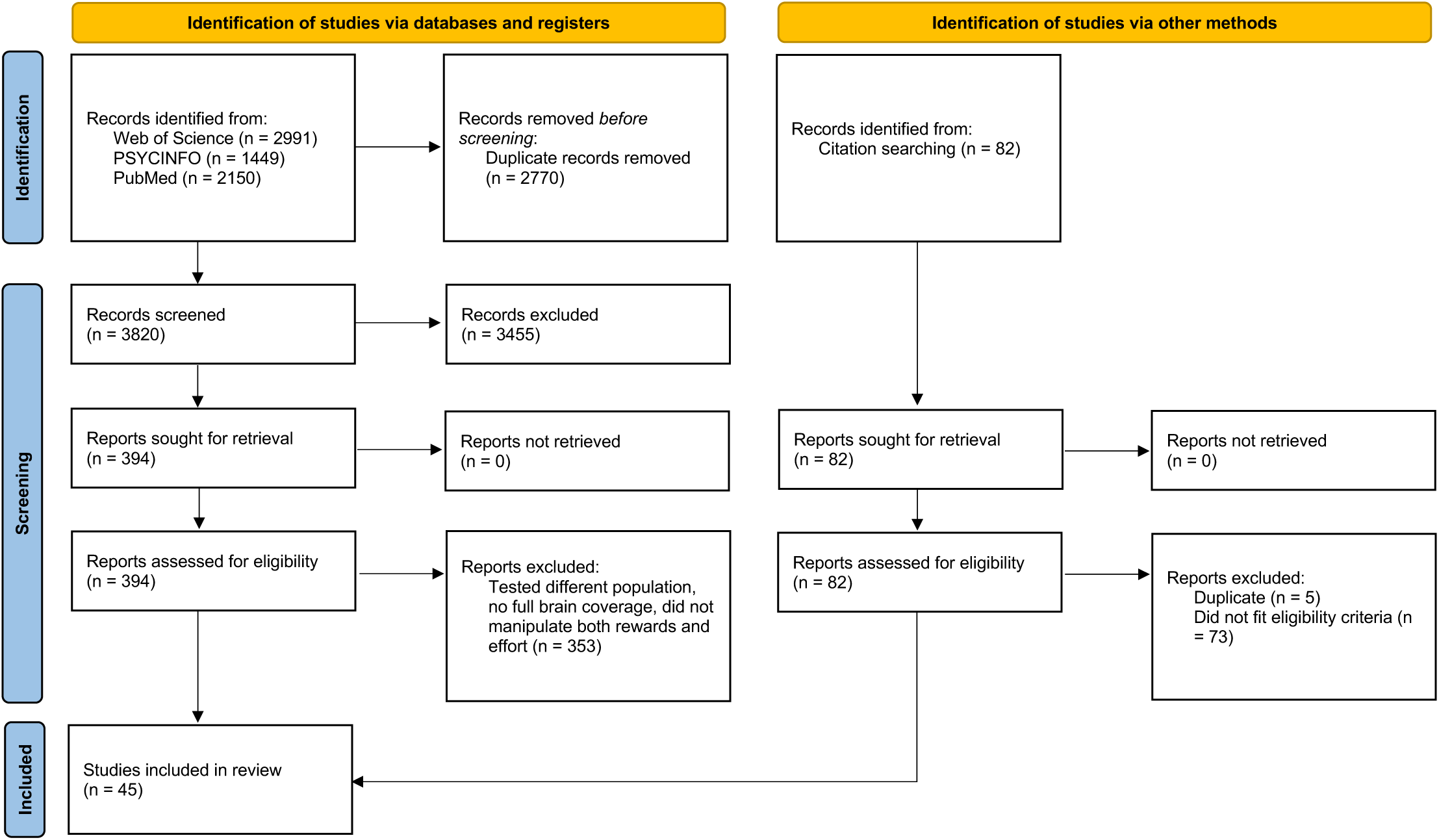
Flowchart of article screening and selection, following PRISMA guidelines (Page et al., 2021).

We screened the identified articles for the following inclusion criteria: 1) empirical investigations (i.e., not review articles); 2) studies employing fMRI; 3) studies carried out in healthy adult samples (i.e. comprised of participants at least 18 years old) excluding studies investigating older adult or individuals with clinical disorders; 4) studies estimate effects using GLMs over the whole brain with reported Montreal Neurologic Institute (MNI) or Talairach coordinates; 5) studies reporting main effects of reward and/or demand level upon fMRI BOLD activity. For the reward manipulations, the reward had to be 1) instrumental (i.e., based on responses) 2) performance-contingent (i.e., not random) 3) mediated by the successful engagement of cognitive processes (e.g., attention, working memory, response inhibition, etc.) as opposed to physical exertion and 4) not serve as a distractor (e.g., Failing & Theeuwes, 2017).

For the mental demand manipulations, demand level had to be manipulated experimentally—note, here we assume increasing task demands require greater effort to resolve (Shenhav et al., 2013). Using these criteria, we included a total of 45 studies (in a total 1,270 participants), with 46 independent samples as one study reported 2 experiments (Ursu et al., 2008). We should note that we obtained two independent sets of contrasts from one study (Kouneiher et al., 2009), and one study collapsed analyses across samples of young adults and adolescents (Magis-Weinberg et al., 2019).

### Coordinate based meta-analysis

We ran a coordinate-based meta-analysis using the foci (i.e., coordinates in significant clusters) reported in the identified studies. To ensure all coordinates were in the same stereotaxic space, we transformed the coordinates reported in Talairach space to MNI space using the FSL transformation applied in GingerALE (Eickhoff et al., 2012). The *x* (left vs right) and *y* (anterior vs posterior) coordinates of one paper was identified as inverted based on the anatomical labels reported (Chikara et al., 2018), and accordingly, we multiplied these coordinates by -1 to convert them back into standard space. We excluded any coordinates identified to be outside the brain, this resulted in the removal of 3 foci (2 from Reward contrasts and 1 from Control contrasts), and a final sample of 429 foci for rewards and 460 foci for task demands (see Table 1). In addition, our literature search revealed 32 foci associated with deactivations for increasing reward prospects, 16 each from 2 studies (Krebs et al., 2012; Pochon et al., 2002) and 8 foci increasing task demands (Krebs et al., 2012). We opted to exclude these coordinates from the analysis given our specific interest in identifying regions encoding task demand and reward incentive levels.

**Table 1.**
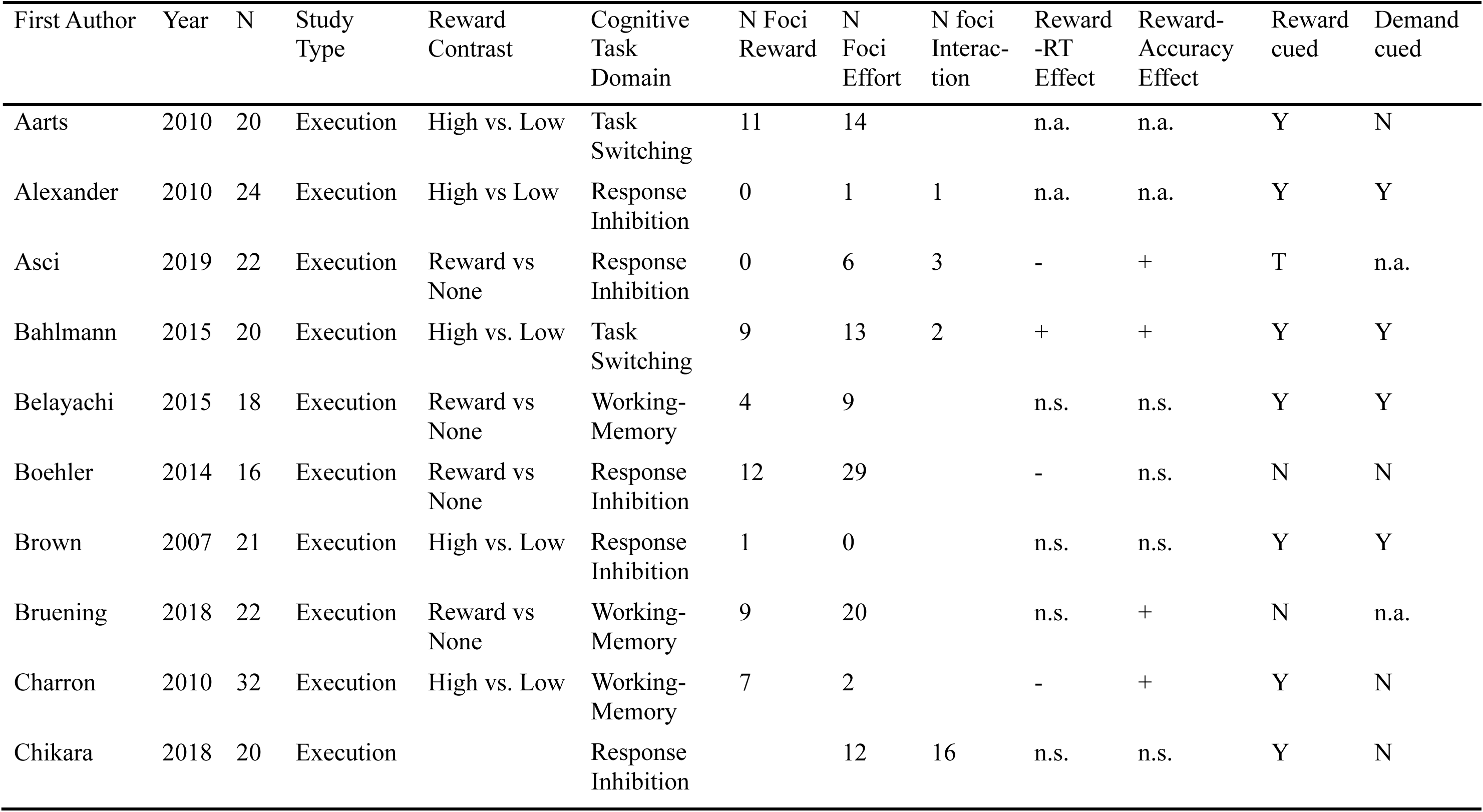
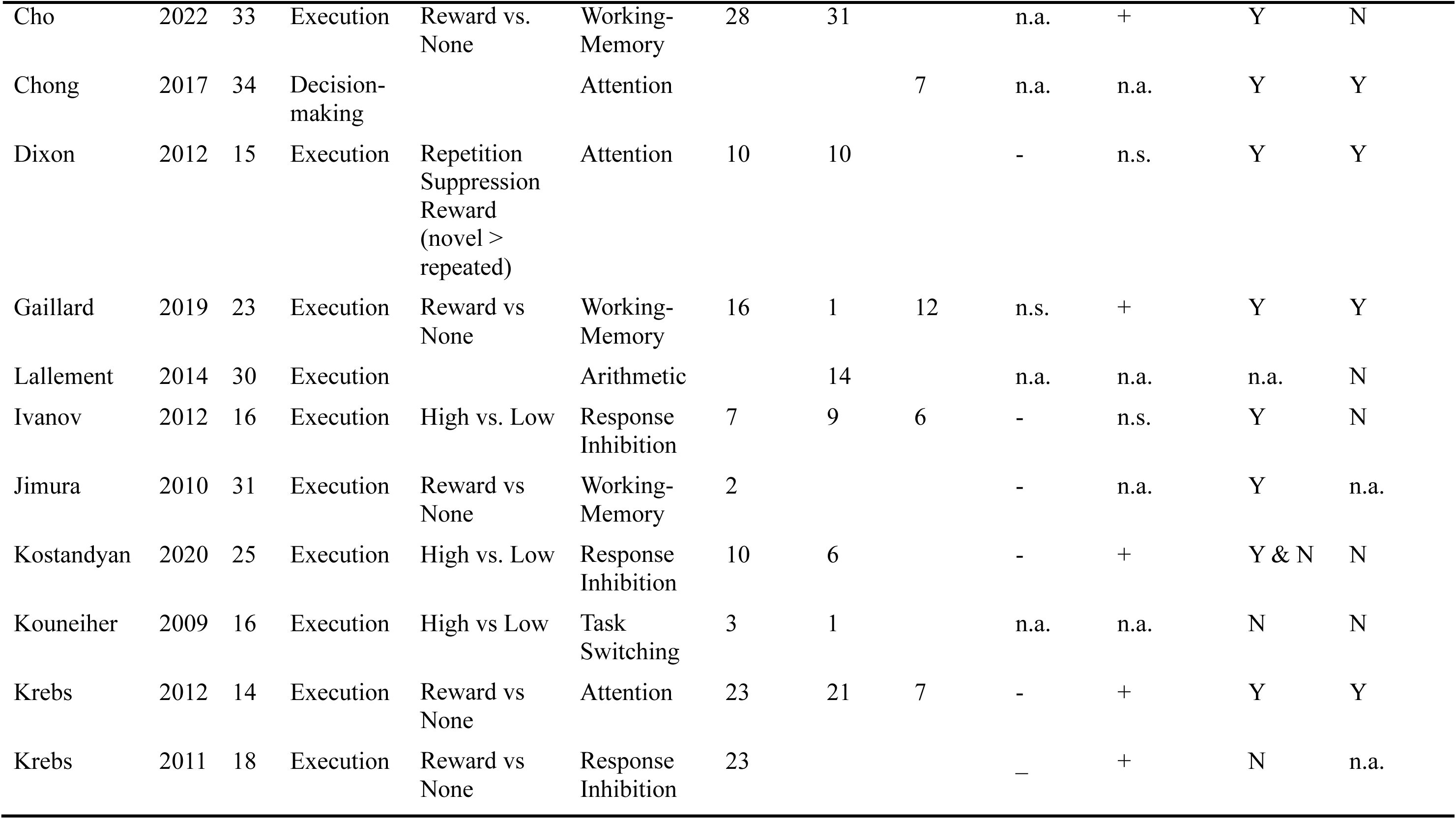
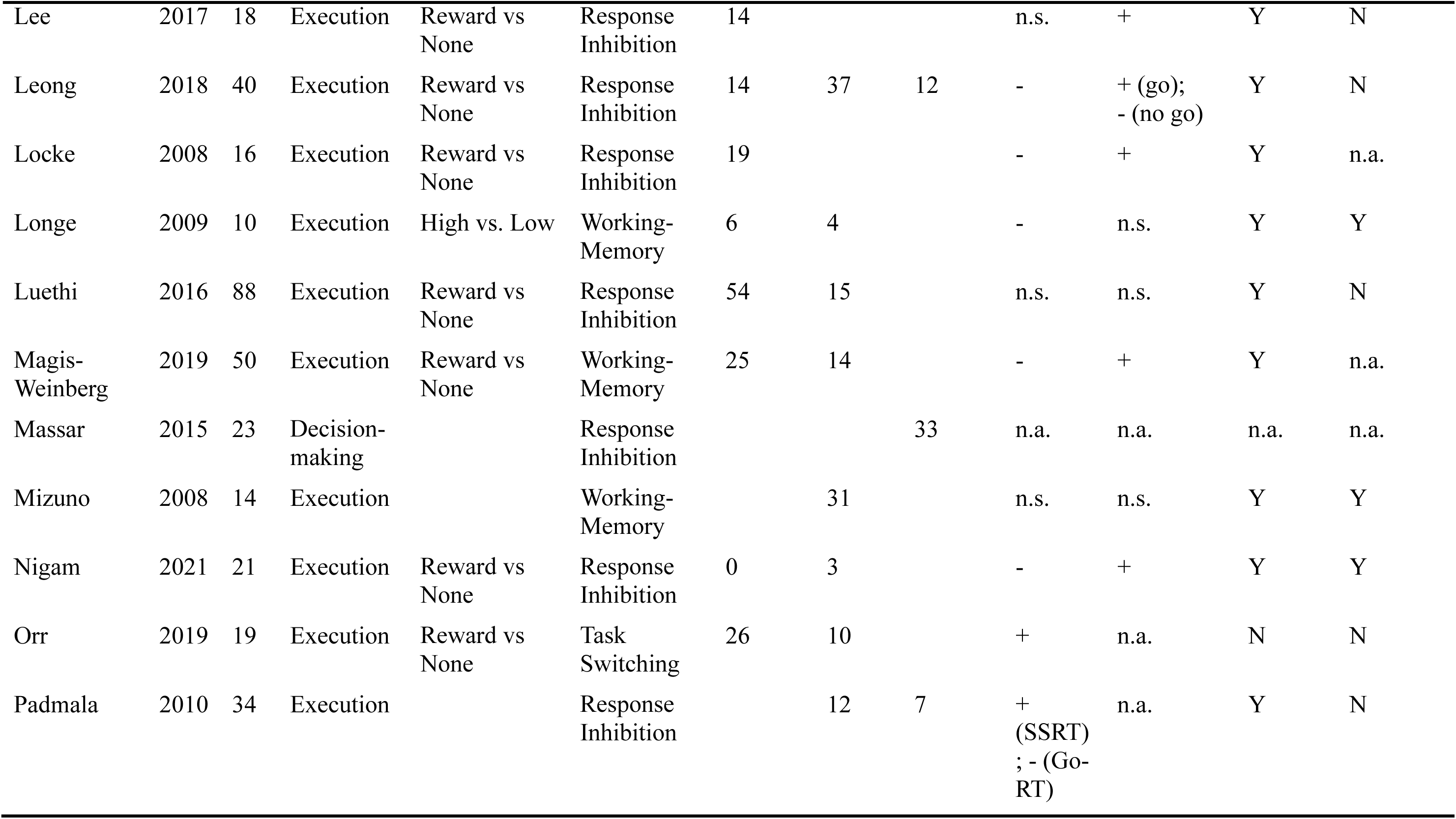
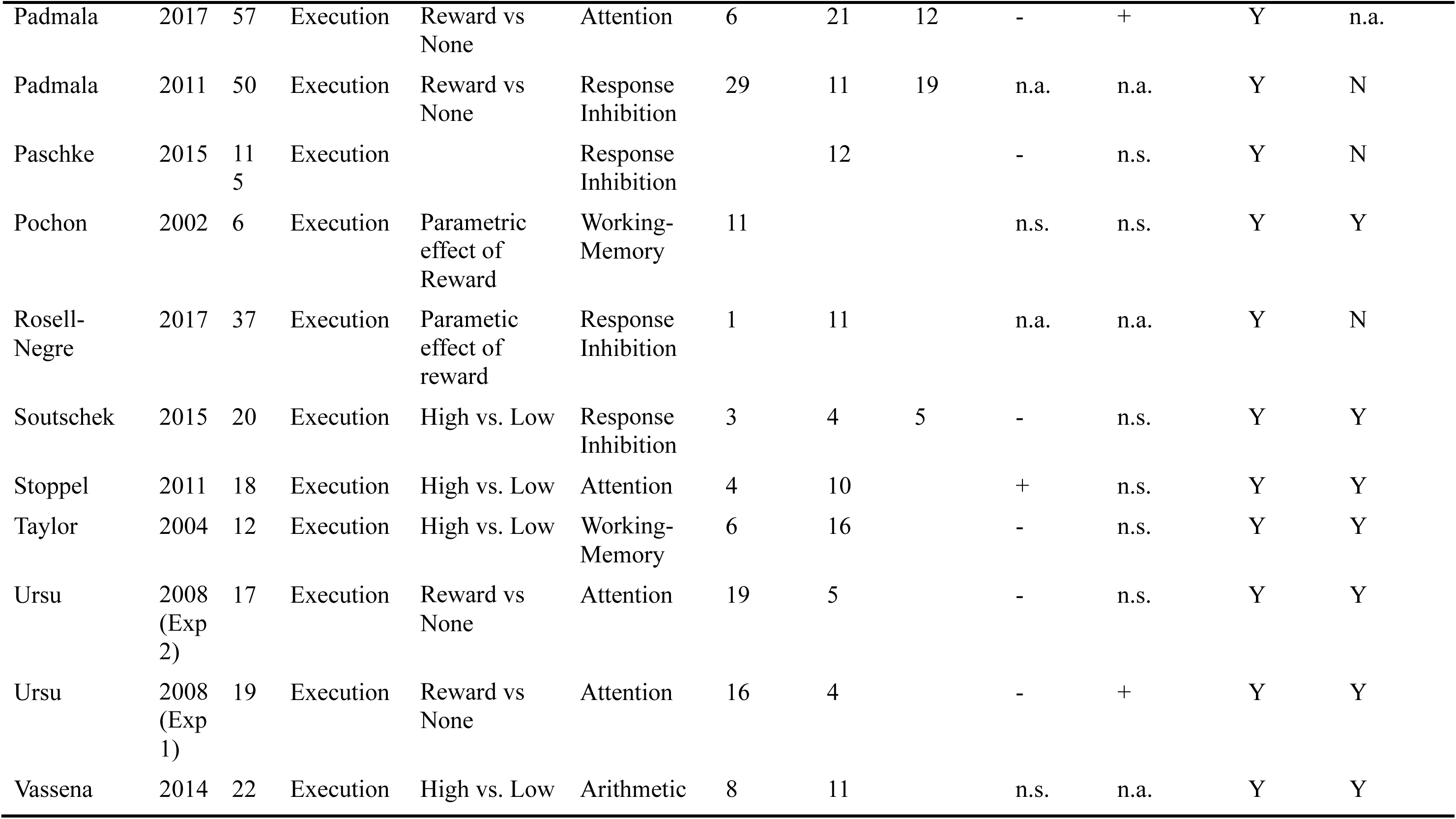
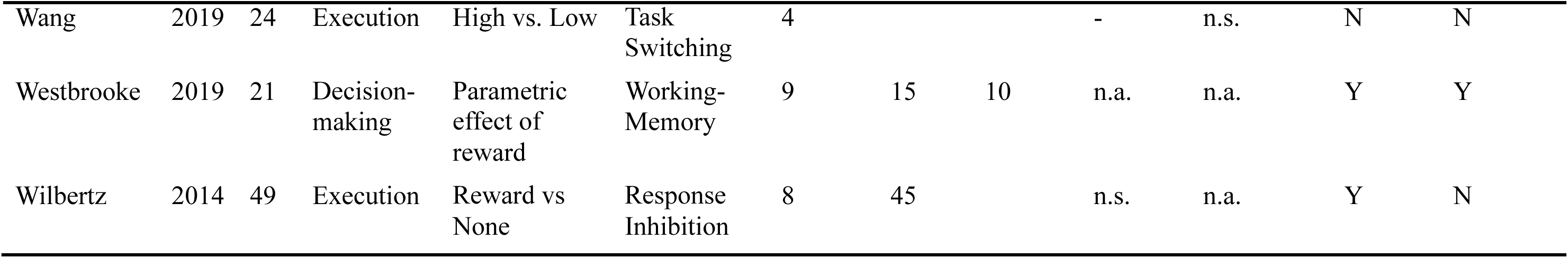
List of reward and effort studies meeting inclusion criteria.

| First Author | Year | N | Study Type | Reward Contrast | Cognitive Task Domain | N Foci Reward | N Foci Effort | N foci Interaction | Reward -RT Effect | Reward-Accuracy Effect | Reward cued | Demand cued |
| --- | --- | --- | --- | --- | --- | --- | --- | --- | --- | --- | --- | --- |
| Aarts | 2010 | 20 | Execution | High vs. Low | Task Switching | 11 | 14 |  | n.a. | n.a. | Y | N |
| Alexander | 2010 | 24 | Execution | High vs Low | Response Inhibition | 0 | 1 | 1 | n.a. | n.a. | Y | Y |
| Asci | 2019 | 22 | Execution | Reward vs None | Response Inhibition | 0 | 6 | 3 | - | + | T | n.a. |
| Bahlmann | 2015 | 20 | Execution | High vs. Low | Task Switching | 9 | 13 | 2 | + | + | Y | Y |
| Belayachi | 2015 | 18 | Execution | Reward vs None | Working-Memory | 4 | 9 |  | n.s. | n.s. | Y | Y |
| Boehler | 2014 | 16 | Execution | Reward vs None | Response Inhibition | 12 | 29 |  | - | n.s. | N | N |
| Brown | 2007 | 21 | Execution | High vs. Low | Response Inhibition | 1 | 0 |  | n.s. | n.s. | Y | Y |
| Bruening | 2018 | 22 | Execution | Reward vs None | Working-Memory | 9 | 20 |  | n.s. | + | N | n.a. |
| Charron | 2010 | 32 | Execution | High vs. Low | Working-Memory | 7 | 2 |  | - | + | Y | N |
| Chikara | 2018 | 20 | Execution |  | Response Inhibition |  | 12 | 16 | n.s. | n.s. | Y | N |
| Cho | 2022 | 33 | Execution | Reward vs. None | Working-Memory | 28 | 31 |  | n.a. | + | Y | N |
| Chong | 2017 | 34 | Decision-making |  | Attention |  |  | 7 | n.a. | n.a. | Y | Y |
| Dixon | 2012 | 15 | Execution | Repetition Suppression Reward (novel > repeated) | Attention | 10 | 10 |  | - | n.s. | Y | Y |
| Gaillard | 2019 | 23 | Execution | Reward vs None | Working-Memory | 16 | 1 | 12 | n.s. | + | Y | Y |
| Lallement | 2014 | 30 | Execution |  | Arithmetic |  | 14 |  | n.a. | n.a. | n.a. | N |
| Ivanov | 2012 | 16 | Execution | High vs. Low | Response Inhibition | 7 | 9 | 6 | - | n.s. | Y | N |
| Jimura | 2010 | 31 | Execution | Reward vs None | Working-Memory | 2 |  |  | - | n.a. | Y | n.a. |
| Kostandyan | 2020 | 25 | Execution | High vs. Low | Response Inhibition | 10 | 6 |  | - | + | Y & N | N |
| Kouneiher | 2009 | 16 | Execution | High vs Low | Task Switching | 3 | 1 |  | n.a. | n.a. | N | N |
| Krebs | 2012 | 14 | Execution | Reward vs None | Attention | 23 | 21 | 7 | - | + | Y | Y |
| Krebs | 2011 | 18 | Execution | Reward vs None | Response Inhibition | 23 |  |  | - | + | N | n.a. |
| Lee | 2017 | 18 | Execution | Reward vs None | Response Inhibition | 14 |  |  | n.s. | + | Y | N |
| Leong | 2018 | 40 | Execution | Reward vs None | Response Inhibition | 14 | 37 | 12 | - | +(go);<br>-(no go) | Y | N |
| Locke | 2008 | 16 | Execution | Reward vs None | Response Inhibition | 19 |  |  | - | + | Y | n.a. |
| Longe | 2009 | 10 | Execution | High vs. Low | Working-Memory | 6 | 4 |  | - | n.s. | Y | Y |
| Luethi | 2016 | 88 | Execution | Reward vs None | Response Inhibition | 54 | 15 |  | n.s. | n.s. | Y | N |
| Magis-Weinberg | 2019 | 50 | Execution | Reward vs None | Working-Memory | 25 | 14 |  | - | + | Y | n.a. |
| Massar | 2015 | 23 | Decision-making |  | Response Inhibition |  |  | 33 | n.a. | n.a. | n.a. | n.a. |
| Mizuno | 2008 | 14 | Execution |  | Working-Memory |  | 31 |  | n.s. | n.s. | Y | Y |
| Nigam | 2021 | 21 | Execution | Reward vs None | Response Inhibition | 0 | 3 |  | - | + | Y | Y |
| Orr | 2019 | 19 | Execution | Reward vs None | Task Switching | 26 | 10 |  | + | n.a. | N | N |
| Padmala | 2010 | 34 | Execution |  | Response Inhibition |  | 12 | 7 | +(SSRT)<br>; -(Go-RT) | n.a. | Y | N |
| Padmala | 2017 | 57 | Execution | Reward vs None | Attention | 6 | 21 | 12 | - | + | Y | n.a. |
| Padmala | 2011 | 50 | Execution | Reward vs None | Response Inhibition | 29 | 11 | 19 | n.a. | n.a. | Y | N |
| Paschke | 2015 | 115 | Execution |  | Response Inhibition |  | 12 |  | - | n.s. | Y | N |
| Pochon | 2002 | 6 | Execution | Parametric effect of Reward | Working-Memory | 11 |  |  | n.s. | n.s. | Y | Y |
| Rosell-Negre | 2017 | 37 | Execution | Parametric effect of reward | Response Inhibition | 1 | 11 |  | n.a. | n.a. | Y | N |
| Soutschek | 2015 | 20 | Execution | High vs. Low | Response Inhibition | 3 | 4 | 5 | - | n.s. | Y | Y |
| Stoppel | 2011 | 18 | Execution | High vs. Low | Attention | 4 | 10 |  | + | n.s. | Y | Y |
| Taylor | 2004 | 12 | Execution | High vs. Low | Working-Memory | 6 | 16 |  | - | n.s. | Y | Y |
| Ursu | 2008 (Exp 2) | 17 | Execution | Reward vs None | Attention | 19 | 5 |  | - | n.s. | Y | Y |
| Ursu | 2008 (Exp 1) | 19 | Execution | Reward vs None | Attention | 16 | 4 |  | - | + | Y | Y |
| Vassena | 2014 | 22 | Execution | High vs. Low | Arithmetic | 8 | 11 |  | n.s. | n.a. | Y | Y |

| First Author | Year | N | Study Type | Reward Contrast | Cognitive Task Domain | N Foci Reward | N Foci Effort | N foci Interaction | Reward -RT Effect | Reward-Accuracy Effect | Reward-cued | Demand-cued |
| --- | --- | --- | --- | --- | --- | --- | --- | --- | --- | --- | --- | --- |
| Wang | 2019 | 24 | Execution | High vs. Low | Task Switching | 4 |  |  | - | n.s. | N | N |
| Westbrooke | 2019 | 21 | Decision-making | Parametric effect of reward | Working-Memory | 9 | 15 | 10 | n.a. | n.a. | Y | Y |
| Wilbertz | 2014 | 49 | Execution | Reward vs None | Response Inhibition | 8 | 45 |  | n.s. | n.a. | Y | N |

We preformed meta-analyses using GingerALE (3.0.2; Eickhoff et al., 2009, 2012; availalble at www.brainmap.org/ale). The Activation Likelihood Estimation (ALE) algorithm computes convergence of activation across coordinates reported from whole-brain analysis. To do so, ALE models the spatial uncertainty of coordinates using 3-dimensional full width at half maximum (FWHM) gaussian kernels centered at the foci, with a width inversely proportional to the sample size. Thus, coordinates from studies with larger sample sizes are modeled with smaller Gaussian kernels, reflecting a more reliable approximation of the true spatial location of BOLD activity. Conversely, coordinates from studies with smaller sample sizes are modeled with larger Gaussian kernels, reflecting the uncertainty in the precise spatial location of activity. Using these activation likelihood estimates, GingerALE computes the overlap of activation probabilities and determines voxels where there is a convergence significantly higher than expected if results were independently distributed. The resulting images can then be corrected for multiple comparisons, using cluster correction. For the purposes of our analysis, we chose a relatively conservative threshold (*p* < 0.05 FWE; 5000 permutations, *p* < 0.001 cluster forming threshold). To characterize the specific cluster location, we use the terms rostral, caudal and dorsal, and we invite the reader to cross-check coordinates when drawing specific conclusions for future studies.

To test whether both rewards and mental demand reliably engage the dACC, we estimated three separate meta-analyses on studies manipulating 1) rewards (36 studies, 920 participants); 2) task demands (38 studies, 1095 participants) and 3) reported interactions between rewards and task demands (15 studies, 418 participants). Additionally, we ran a conjunction/contrast analyses comparing reward to effort. Conjunction between two sets of coordinates can be assessed using the voxel-wise minimum value of the activation likelihood estimates (Eickhoff et al., 2012). Contrasting the two sets of coordinates is done by subtracting the activation likelihood estimates between images and calculating voxel-wise Z-scores of the differences against a permuted distribution (Eickhoff et al., 2012). These resulting Z-scored differences are then subject to cluster analysis. For our contrast analysis, we conducted 100,000 permutations and set a threshold *p* < 0.01 FWE and minimum cluster size of 300mm^3^.

Given the heterogeneity of the studies included in this meta-analysis with respect to their operationalization of task demands and incentives, statistical power is an important consideration. Current best practices suggest the inclusion of a minimum of 17-20 studies in a meta-analysis to ensure sufficient power and mitigate the possibility of results being driven by a single study (Eickhoff et al., 2016; Müller et al., 2018). Our literature search revealed 45 studies (46 total experiments), reporting a total of 36 reward contrasts and 38 demand contrasts of interest, resulting in 457 reward-related foci and 491 demand-related foci (see Table 1). Of the 46 studies included, 34 reported both reward and effort contrasts. A total of three reward contrasts, and one demand contrasts observed no significant foci for their contrast of interest. With respect to reward contrasts, 21 studies contrasted reward with no reward, while 14 studies reported a contrast between different reward levels, 3 studies reported a parametric effect of reward, 1 study used a repetition suppression paradigm (see Table 1). With respect to effort contrasts, the most common operationalization of task demand involved response inhibition (20 studies), followed by working memory (12 studies), attention (7 studies), task-switching (6 studies), and arithmetic (2 studies; see Table 1). Critically, of the experiments which investigated reward effects on task performance (45 experiments), 9 examined accuracy differences, 16 tested response time differences, 15 tested both measures, and 3 employed effort discounting choice paradigms. Most studies reported significant improvements in task performance with increasing rewards (36 of 45 studies; see Table 1).

## Results

### BOLD response to rewards

First, we sought to test which brain regions reliably encoded information about performance-contingent rewards. To this end, we assessed patterns of brain activity during task execution that reliably differentiated between reward incentive levels, observing six clusters that were sensitive to higher reward prospect. These clusters were located the bilateral putamen, as well as the dorsal portion of the mPFC/dACC. We also found activation in the right anterior insula hand left inferior occipital cortex (see Table S1 for a complete list of clusters and MNI coordinates).

### BOLD response to task demand

Next, we sought to test which brain areas display increased as a function of task demand level during task execution (see Table S2 for a complete list of clusters and MNI coordinates). Our analysis revealed a reliable pattern of increasing brain activity associated with increased mental demand in several clusters across regions typically associated with cognitive control (Laird et al., 2005; Niendam et al., 2012) including mPFC/dACC, lateral PFC, parietal cortex and bilateral insula. Specifically, we observed increased activity for higher demand in the dorsal portion of the mPFC/dACC (extending to the pre-SMA). Activation clusters were also observed in the left middle frontal gyrus (extending to the left precentral gyrus) and in the right inferior frontal gyrus (extending posteriorly to the right precentral gyrus). Additionally, we observed bilateral activation of the superior parietal lobule extending into the precuneus, and a cluster in the right thalamus.

Subsequently, we sought to examine whether activations differed across studies where task demand was cued versus uncued (prior to task execution). To this end, we conducted a follow-up analysis directly contrasting BOLD responses across these two study types (i.e., cued demand versus uncued demand; see Figure S1 of the Supplemental Materials). This analysis revealed one significant cluster resulting from the Cued > Uncued demand level contrast in the dorsal portion of the mPFC/dACC, and one significant cluster resulting from the Uncued > Cued demand level contrast in the inferior frontal gyrus (see Table S3 for cluster details). It is important to note that only 19 studies could be used for this comparison, and the results should be interpreted with caution due to lower power.

### Differences in BOLD response between reward versus task demand

Next, we tested unique and overlapping activity between reward and mental demand-supporting task demand level to address the key question of whether mPFC/dACC activity tracks mainly task demand (Loperz Gamundi et al., 2021) or reward and demand integration (Chong et al, 2017, Shenhav et al. 2013, Silvetti et al. 2018) (see Figure 1). For larger rewards (compared to demands) we observed one cluster in the rostral portion of the mPFC/dACC was more active (see Table 2, and yellow cluster depicted in Figure 3). Conversely, for higher rewards (compared to demands) we observed six clusters that were more active, including a bilateral cluster in the dorsal portion of the mPFC/dACC extending into the pre-SMA, as well as two clusters in the left middle frontal gyrus, the bilateral inferior parietal lobule, and the right precuneus (see Table 2, and blue clusters depicted in Figure 3). Finally, we tested the overlap between demand and reward, revealing regions that were more active for both higher demand levels and larger reward incentives, observing a cluster in the right anterior insular cortex (see Table 2, and green clusters depicted in Figure 3). Together, these results highlight a distinction between mPFC/dACC responses to rewards and task demands: task demand was associated with activity in in the dorsal portion of dACC, extending into the pre-SMA, whereas reward was associated with activity in the rostral portion of the mPFC/dACC.

**Figure 3.**
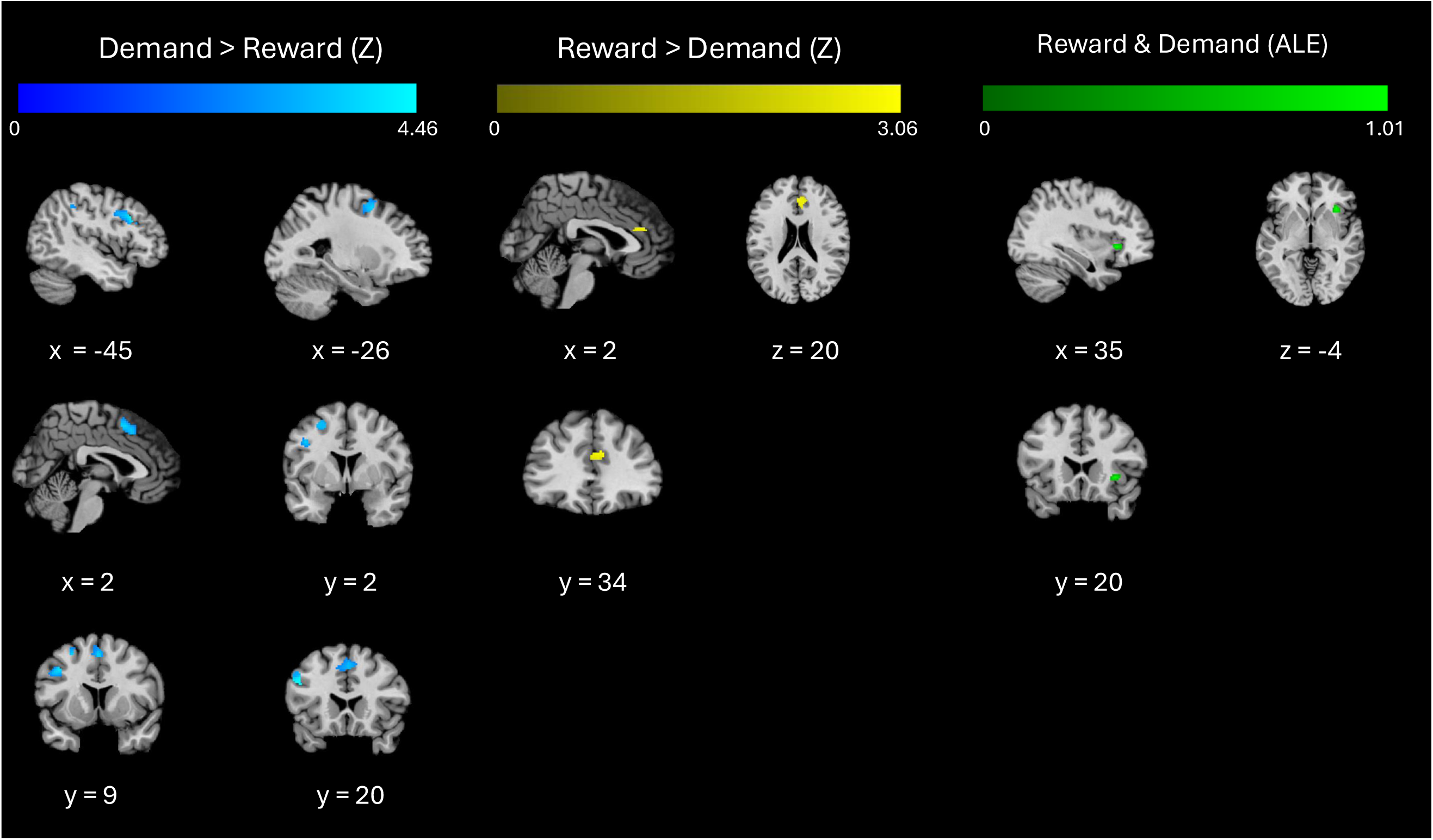
Results of reward and effort ALE meta-analysis. Left Columm: Brain areas activated more by increasing task demands than by reward prospects are plotted with shades of blue . Middle Column: Brain areas actived more by increasing reward prospects than task demands are plotted with shades of yellow. Right Column: Brain areas exhibiting converging activation for both increasing reward prospects and task demands are plotted with shades of green.

**Table 2.**
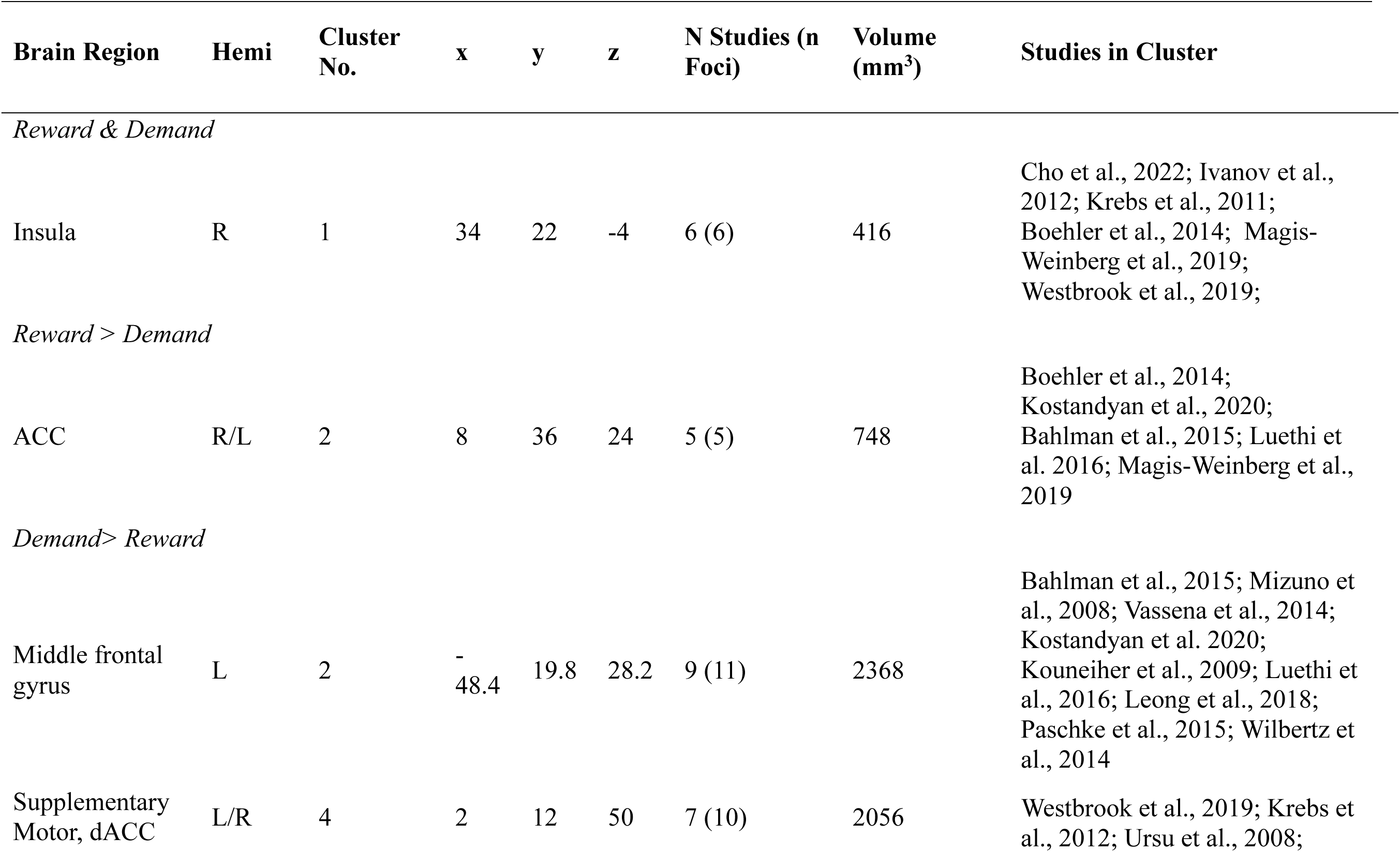
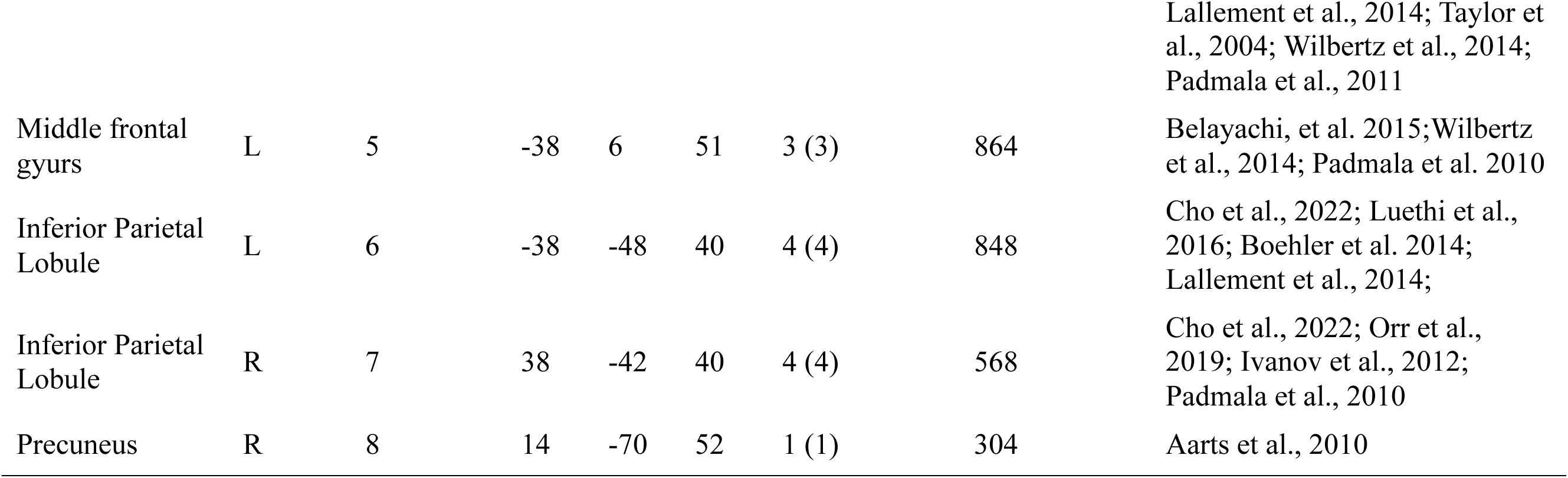
Conjunction and difference of the ALE meta-analysis for demand and reward. For each coordinate, region label, hemisphere (right, left or bilateral), Brodmann area, MNI coordinates, ALE maxima, *p* values, *Z* values, number of studies, and cluster sizes are provided.

### BOLD response to integrated cost-benefits

Next, we explored whether there were any reliable patterns of activation associated with the integration of effort and rewards across 15 experiments (152 foci from 418 participants)—as indexed by BOLD response correlated with either computed subjective value in effort discounting tasks (Chong et al., 2017; Massar et al., 2015; Westbrook et al., 2019) or by interactions between reward and task demands in reward-incentivized control studies. Our analysis revealed one cluster in the mPFC/dACC (including the pre-SMA) and one cluster in the middle frontal gyrus (see Table 3, depicted in warm colors in Figure 4). These two clusters partially overlapped with clusters previously identified to respond more to increasing demands versus rewards (i.e., the Demand > Reward contrast; depicted in cool colors in Figure 4). Interesting, the mPFC/dACC cluster contained coordinates from four studies (Bahlmann et al., 2015; Padmala & Pessoa, 2011; Westbrook et al., 2019), three of which reported BOLD responses consistent with cost-benefit integration (see Figure 1 right panel). The middle frontal gyrus cluster contained coordinates from four studies (Chong et al., 2017; Leong et al., 2018; Padmala & Pessoa, 2010, 2011), two of which reported BOLD responses consistent with cost-benefit integration (see Figure 1, right panel). Taken together, these results support the hypothesis that mPFC/dACC activity subserves integration of mental demand and expected rewards, compatible with functional accounts positing a cost-benefit trade-off.

**Figure 4.**
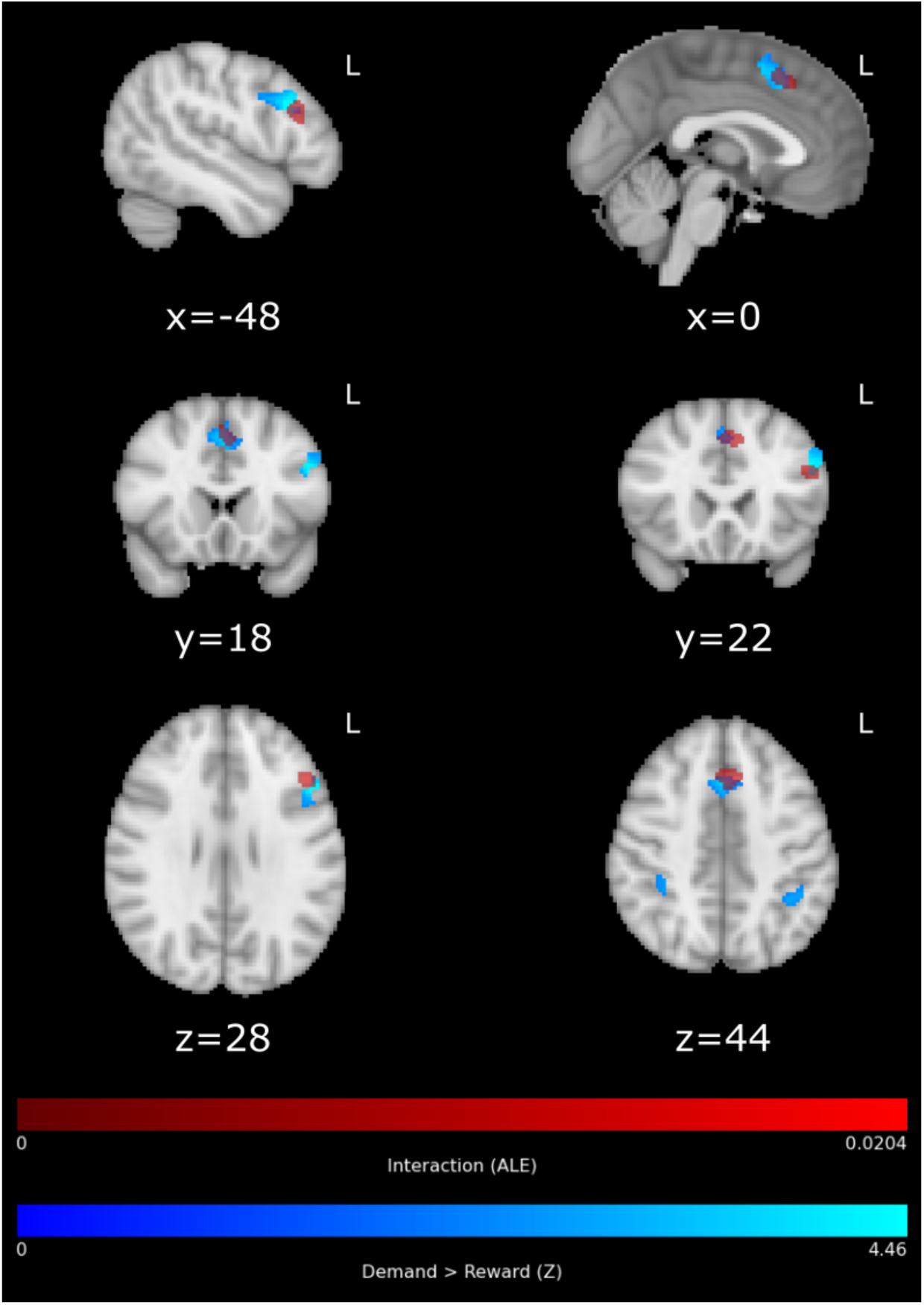
Results of interaction (reward X effort) ALE meta-analysis. Brain areas showing converging activation for the interaction between rewards and effort plotted in the volume with shades of red. Brain areas activated by task demands plotted in the volume with shades of blue. Blue clusters were rendered transparent to depict the overlap between clusters.

**Table 3.**
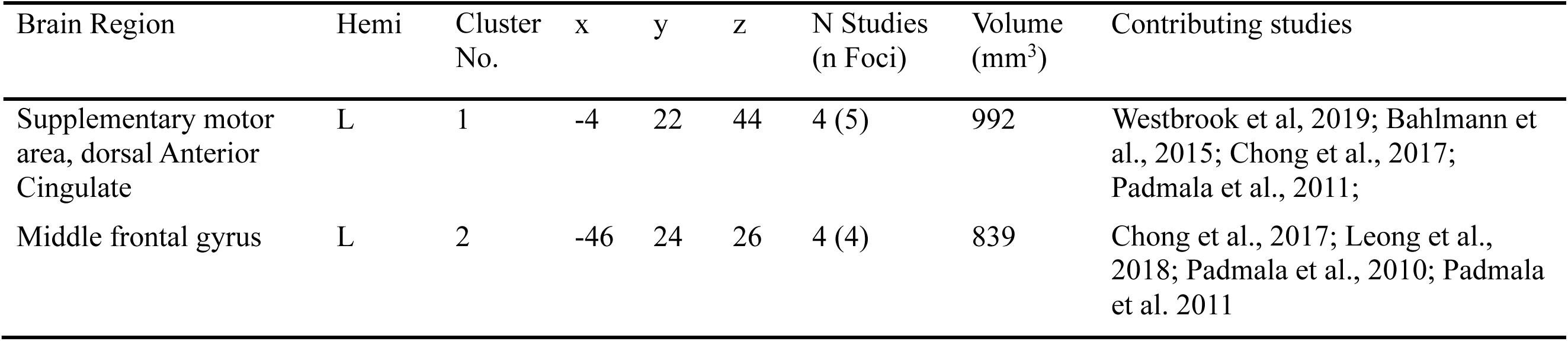
The ALE meta-analysis for coordinates representing the interaction between reward and demand level. For each cluster, region label, hemisphere (right, left or bilateral), the number of studies, and cluster size are provided.

## Discussion

Recent neurocomputational accounts of effort-based decision-making posit that activity in the medial prefrontal cortex, and particularly the dACC (mPFC/dACC) reflects a trade-off between the costs and benefits of allocating mental effort (Silvetti et al., 2018; Verguts et al., 2015). Yet, the precise functional role of the dACC in effortful behavior remains debated (Vassena et al., 2017, 2020, Vriens et al. 2025). Prior meta-analytic work—including studies on both physical and mental demand —concluded that a dorsal portion of mPFC/dACC uniquely tracks task-demand, rather than a trade-off per se (Lopez-Gamundi et al., 2021)we carried out a meta-analysis of neuroimaging studies jointly manipulating mental demand and reward incentives, and assessed whether mPFC/dACC activity reflected reward, mental demand, or an integration of both, compatible with a cost-benefit signal. Our results showed that increasing task demand was associated with activity in the mPFc/dACC, insular, frontal and parietal cortices—replicating extant work (Laird et al., 2005; Niendam et al., 2012). Increased reward prospects engaged the rostral portion of the mPFC/dACC. However, the only overlap observed for rewards and demand was located in the Insula, with no overlap in the dorsal mPFC/dACC. Instead, we found two dissociable cluster within dACC/mPFC — the dorsal portion tracking mental demand, and the rostral portion tracking reward incentives, supporting the view that effort costs and rewards are represented separately in the brain. Activity in the dorsal portion of the mPFC/dACC activity also tracked the interaction between mental demand and rewards providing evidence for an integrative cost-benefit account (Chong et al., 2017; Shenhav et al., 2013; Silvetti et al., 2018; Soutschek & Tobler, 2020). These results bring together prior and seemingly conflicting findings: rostral mPFC/dACC and dorsal mPFC/dACC uniquely tracked rewards and mental demands respectively, while dorsal mPFC/dACC also tracked cost-benefit integration.

The functional role of mPFC/dACC has been highly debated (Ebitz & Hayden, 2016; Vassena et al., 2017). Our results provide a clearer understanding of the mPFC/dACC’s role in motivating deployment of mental effort. Prominent theories of cognitive effort support cost-benefit models where the control signal intensity is determined by the mPFC/dACC, which integrates both information about available rewards and the cost associated to exerting control (Shenhav et al., 2013, 2016). In line with previous findings, the present results provide indication for a role of the mPFC/dACC in tracking task demand (Lopez-Gamundi et al., 2021) as well a signal integrating demand level and reward (Chong et al., 2017; Shenhav et al., 2013, 2016; Silvetti et al., 2018). 4), We did not observe overlap in the mPFC/dACC in the tracking of demands and rewards, in contrast to the findings of a number of prior studies included in this meta-analysis (Boehler et al., 2014; Krebs et al., 2012; Paschke et al., 2015; Vassena et al., 2014). It is possible that this previously observed overlap may have been driven by task-specific factors—for example, obtaining larger rewards in that task may have elicited subjectively larger efforts (despite objective demand being controlled across reward conditions). Plausibly, activity in the mPFC/dACC may scale with reward-incentivez efforts, rather than rewards per se.

The likelihood of (successful) effort exertion resulting in reward (often termed ‘efficacy’) has indeed been related to mPFC/ACC activity (Frömer et al., 2021; Grahek et al., 2022). At the same time, previous work also identified a role for the mPFC/dACC in monitoring the need for cognitive control (Botvinick, 2007; Venkatraman & Huettel, 2012) but also effort-discounted rewards (Chong et al., 2017). The results of our meta-analysis converge with his line of evidence, suggesting that the dorsal portion of mPFC/dACC tracks the integrated cost-benefit signal. Interestingly, this location overlapped with the portion of mPFC/dACC tracking increasing mental demands. This overlap may reflect a monotonic relationship between mental demands and effort investment (the harder the task, the more effort invested), which is particularly realistic where most of the tasks manipulating mental demands are titrated to obtain (relatively) high performance accuracy. In other words, these tasks are difficult but still feasible. However, this monotonic relationship may not always apply, for example when greater effort does not yield better performance, because the task is impossible (Gendolla et al., 2019; Otto et al., 2022; Silvestrini et al., 2022), in line with the well-characterized inverted-U shaped relationships between task difficulty and effort investment, where mental effort investment has been observed to decrease under particularly high demand levels (Bognar et al., 2024; da Silva Castanheira & Otto, in press).

Neurally, a body of neurocomputational work suggests that the dorsal portion of mPFC/dACC contributes to learning optimal effort allocation through reinforcement meta-learning (Silvetti et al., 2018; Verguts et al., 2015) and by tracking negative feedback (i.e., response errors) (Carter et al., 1998; Cole et al., 2009; Ito et al., 2003; Ridderinkhof et al., 2004).

On this view, cost-benefit models of dorsal mPFC/dACC function predict that activity should vary depending on the response efficacy (Frömer et al., 2021; Shenhav et al., 2013). When the task is feasible at high demand levels, dorsal mPFC/dACC activity should grow monotonically with task demands. When the task is impossible (i.e., high demand), activity should follow an inverted-U pattern, with a drop in engagement when demand is too high (Silvestrini et al. 2021). Together, these predictions explain how increases in task demand—which purportedly decrease the net value of effort—were associated with both increases and decreases in mPFC/dACC activity in the literature. Beyond dACC activity, other physiological measures like cardiovascular reactivity (see Wright, 2008 for discussion), pupil dilation (da Silva Castanheira et al., 2021), and facial muscle activity (de Morree & Marcora, 2010; Devine et al., 2023; Van Boxtel & Jessurun, 1993) have been proposed as a method for indirectly measuring effort exertion. To reconcile these findings regarding the functional role of the mPFC/dACC in effortful behavior, future work should approach triangulation by jointly considering rewards, task demands and efficacy alongside neural activity, and psychophysiological measures of effort exertion.

Notably, the studies examined in this meta-analysis were constrained to those which used monetary incentives to motivate mental effort. Beyond cognitive control, previous work has found evidence for the mPFC/dACC’s involvement in processing and integrating both primary and secondary rewards (Yee et al., 2021). Similarly, dorsal mPFC/dACC has been found to integrate information about physical effort (Chong et al., 2017), pain and negative affect (Shackman et al., 2011). While cost-benefit models posit that the dorsal mPFC/dACC integrates signals reflecting general costs and benefits, more work is needed to understand whether this pattern generalizes to other stimuli—particularly as there is some evidence for an anterior-posterior gradient of functional specialization from strategic to response-related conflict (Alexander & Brown, 2015; Venkatraman et al., 2009). The results of our reward-demand contrast analysis—in which rostral mPFC/dACC was found to respond more reliably to rewards while dorsal mPFC/dACC responded to demand—coincides with previous work which also suggests functional specialization of the mPFC/dACC: a cognitive-affective gradient moving from caudal to rostral dACC (Bush et al., 2000). However, these distinctions have been inconsistent as others have found cognitive demand, affect and pain to overlap in the same region (Shackman et al., 2011). To what extent the rostral portion of MPFC/dACC uniquely responds to value, or possibly to unsigned incentives or salience, remains to clarified. At the same time, while much of the existing work examining effort-reward decision-making (tacitly) assume that mental demand level and reward are separate (or even orthogonal) quantities, it is well documented that some individuals find effort exertion to be intrinsically rewarding (Inzlicht et al., 2018; Job et al., 2024), which may pose conceptual challenge for neuroscientific studies endeavoring to understand effort costs separately from extrinsic rewards.

It is also worth noting that this meta-analysis—by design—examined a heterogeneous body of mental effort studies which employed diverse demand and reward incentive manipulations. While there was considerable variability in how these study designs instantiated (and modeled) these factors, we should note that this inherent source of variability may have constrained our ability to detect consistent activity patterns across studies. On the one hand, this across-study variability could be seen as an inherent limitation of the meta-analytic approaches more generally. On the other hand, that our analysis was able to identify brain regions reliably tracking demand, reward (and their interaction) across studies employing heterogenous designs highlights the potential generality of these regions’ involvement in cost-benefit effort evaluation—for example, mental demand level reliably correlated with dorsal mPFC/dACC activation across diverse operationalizations of mental demand. It is also worth noting a potential limitation stemming from the small body of findings (four studies) informing the observed interaction between reward and demand level in the mPFC/dACC (Table 3). This paucity of studies uncovering such an interaction indicates the need for further studies jointly examining mental demand and reward. A larger body of studies examining such an interaction could enrich the input to future meta-analyses of this sort, further supporting (or refining) the putative role of the mPFC/dACC in cost-benefit integration.

The results of this meta-analysis inform our understanding of the broader involvement of this circuitry in value-based choice more generally. While we specifically examined studies involving mental effort, a considerable number of studies have examined the neural correlates of demand level and reward in the domain of physical effort (cf. meta-analysis by Lopez-Gamundi et al., 2021). For example, two recent studies examining effort-reward trade-offs—whereby participants made choices to engage in greater versus less physical effort for larger versus smaller reward amounts—observed that the relative value of the chosen option was tracked by the mPFC/dACC (Chong et al., 2017; Hogan et al., 2020) and right anterior insula (Hogan et al., 2020). Taken together with the present results—which highlights a role for both of these regions in the integration of mental demand and rewards—these findings suggest a common neural representation and possibly a shared underlying cost-benefit computation for both physical and mental effort (in line with what observed by Chong et al. (2017).

Considering the literature on subjective valuation of actions more generally, a large body of work indicates that the vmPFC (among other regions, including the ventral striatum) underpins a domain-general valuation system (Bartra et al., 2013). Similarly, the recent meta-analysis by Lopez-Gamundi et al. (2021)—which examined both mental and physical effort—found that only the vmPFC reliably and positively tracked the net value of effort. In contrast, we observed that the interaction between mental demand and reward appeared was (positively) tracked by the rostral mPFC/dACC (while a more rostral area of dACC tracked reward incentives alone), suggesting that these integrated cost-benefit computations concerning mental demand may differ in nature from the more rostrally-located signals previously implicated in domain-general valuation (e.g. in the vmPFC).

Effort-related decision-making is thought to rely on the coordinated activity between several brain regions (Ullsperger et al., 2014). While the literature as well as theoretical developments have strongly focused on the mPFC/ACC, empirical evidence indicates that other regions may be sensitive to both manipulations of reward and effort and can play a role in resolving cost-benefit trade-offs. For example, the lateral PFC (LPFC) is involved in maintaining task relevant information in working-memory (Braver, 2012; Burgess & Braver, 2010) and executing cognitive control more generally (Miller & Cohen, 2001). In this study, we observed LPFC for both higher mental demand and integration of mental demand and reward. In line with these results, recent work suggests the LPFC encodes an individual’s capacity to successfully meet task demands, which in turn may signal the likelihood of receiving rewards (Soutschek & Tobler, 2020). At the same time the anterior insula, which is commonly co-activated with the mPFC/dACC (Bartra et al., 2013; Diekhof et al., 2012; Parro et al., 2018), is also implicated in monitoring the need for control (Shenhav et al., 2016). Supporting this view, we found that the anterior insula was reliably engaged by both increasing task demands and reward prospects—possibly suggesting a broader role of the region in processing salient events (i.e., arousal; Uddin, 2015) and subjective awareness (Craig, 2002), both of which are foundational to effort allocation. Together, these results suggest a role for the anterior insula in monitoring one’s current state and detecting changes in the need for control (Nelson et al., 2010) and a role in cognitive processes more generally (Uddin et al., 2014). Previous work has shown a negative coupling between reward-related processing in the ventral striatum and dorsal mPFC/dACC activation (Botvinick et al., 2009). Together with the literature, our results support the notion that the dorsal mPFC/dACC, alongside a coordinated set of other brain regions, is involved in the integration of effort costs (i.e. task demand level) and the benefits conferred by rewards.

However, as ALE-coordinate based analyses of this sort preclude examination of network connectivity between regions, these results are limited in their ability to speak to coordination between regions. Thus, future empirical work should aim to characterize the underlying network dynamics informing effort-based decision-making.

## Open Practices Statement

This manuscript has no associated data.

