## Supplemental Materials for "The neural basis of cost-benefit trade-offs in effort investment: a quantitative activation likelihood estimation meta-analysis"

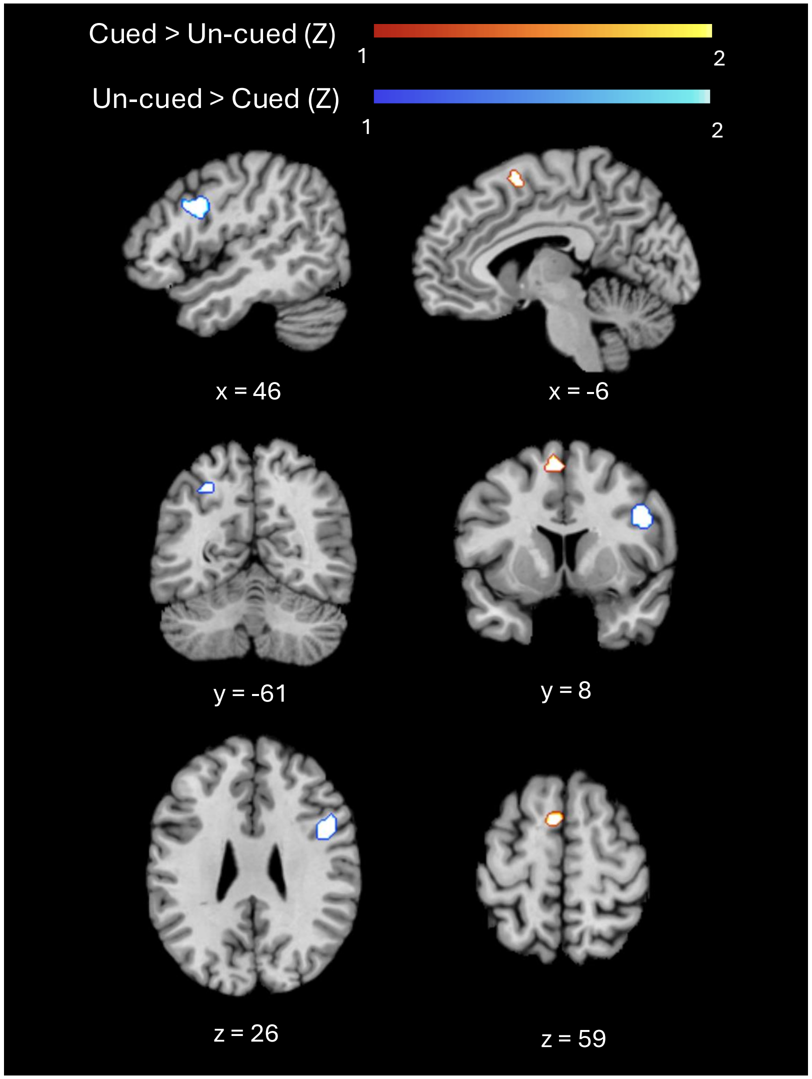


**Figure S1.** Direct contrast comparing activations between Cued and Un-cued Demand levels. The Cued > Un-cued contrast is represented by warm colors, and the Un-cued > Cued contrast is represented by cool colors.

**Table S1.** ALE meta-analysis of clusters tracking Reward level. For each coordinate, region label, hemisphere, Brodmann area (BA), MNI coordinates, ALE maxima, p values, Z values, number of studies, and cluster sizes are provided.

| **Brain Region** | **Hemi** | **BA** | **Cluster No.** | **x** | **y** | **z** | **N Studies (n Foci)** | **Volume ($mm^3$)** | **Studies in Cluster** |
|---|---|---|---|---|---|---|---|---|---|
| *Reward* | | | | | | | | | |
| Putamen | L | | 1 | -10 | 10 | 0 | 11 (16) | 2872 | Westbrook et al., 2019; Krebs et al., 2011; Ursu et al., 2008; Boehler et al., 2014; Kostandyan et al., 2020; Aarts et al., 2010; Dixon & Christoff, 2012; Krebs et al., 2012; Vassena et al., 2014; Luethi et al., 2016; Padmala et al., 2011 |
| Putamen | R | | 2 | 10 | 10 | -2 | 11 (13) | 2472 | Westbrook et al., 2019; Orr et al., 2019; Boehler et al., 2014; Kostandyan et al., 2020; Wang et al., 2019; Aarts et al., 2010; Dixon & Christoff, 2012; Krebs et al., 2012; Leong et al., 2018; Luethi et al., 2016; Padmala et al., 2011 |
| Insula | R | 13 | 3 | 34 | 20 | -6 | 7 (7) | 1320 | Cho et al., 2022; Krebs et al., 2011; Boehler et al., 2014; Kostandyan et al., 2020; Wilbertz et al., 2014; Bahlman |

| | | | | | | | | | |
|---|---|---|---|---|---|---|---|---|---|
| | | | | | | | | | et al., 2015; Magis-Weinberg et al., 2019 |
| Middle Frontal Gyrus | R | 10 | 4 | 42 | 52 | 12 | 7 (9) | 1200 | Charron & Koechlin, 2010; Ursu et al., 2008; Kostandyan et al., 2020; Ivanov et al., 2012; Luethi et al., 2016; Magis-Weinberg et al., 2019; Padmala et al., 2011 |
| Anterior Cingulate Cortex | R/L | 32 | 5 | 8 | 36 | 20 | 7 (8) | 1160 | Boehler et al., 2014; Kostandyan et al., 2020; Bahlman et al., 2015; Dixon & Christoff, 2012; Krebs et al., 2012; Luethi et al., 2016; Magis-Weinberg et al., 2019 |
| Inferior Occipital Gyrus | L | 18 | 6 | -26 | 90 | 2 | 6 (8) | 880 | Cho et al., 2022; Orr et al., 2019; Aarts et al., 2010; Bruening et al., 2018; Magis-Weinberg et al., 2019; Padmala et al., 2011 |

**Table S2.** ALE meta-analysis of clusters tracking Demand level. For each coordinate, region label, hemisphere, Brodmann area (BA), MNI coordinates, ALE maxima, p values, Z values, number of studies, and cluster sizes are provided.

| **Brain Region** | **Hemi** | **BA** | **Cluster No.** | **x** | **y** | **z** | **N Studies (n Foci)** | **Volume (mm$^3$)** | **Studies in Cluster** |
|---|---|---|---|---|---|---|---|---|---|
| Inferior & Superior Parietal Lobules | L | 40 / 7 | 1 | -40 | -42 | 42 | 19 (26) | 5080 | Cho et al., 2022; Bahlman et al., 2015; Mizuno et al., 2008; Belayachi, et al. 2015; Bruening et al., 2018; Krebs et al., 2012; Orr et al., 2019; Soutschek et al., 2015; Kostandyan et al., 2020; Luethi et al., 2016; Boehler et al., 2014; Ivanov et al., 2012; Lallement et al., 2014; Leong et al., 2018; Paschke et al., 2015; Padmala et al., 2010; Padmala et al., 2011; Padmala et al., 2017; Wilbertz et al., 2014 |
| Medial Frontal and Cingulate Gyrus | L/R | 6 / 32 | 2 | -2 | 16 | 46 | 19 (26) | 4992 | Cho et al., 2022; Bahlman et al., 2015; Westbrook et al., 2019; Aarts et al., 2010; Bruening et al., 2018; Dixon & Christoff, 2012; Krebs et al., 2012; Orr et al., 2019; Soutschek et al., 2015; Ursu et al., 2008; Vassena et al., 2014; Kostandyan et al., 2020; Boehler et al., 2014; Ivanov et al., 2012; Lallement et al., 2014; Lallement et al., 2014; Paschke |

| | | | | | | | | | |
|---|---|---|---|---|---|---|---|---|---|
| | | | | | | | | | et al., 2015; Taylor et al., 2004; Wilbertz et al., 2014; Padmala et al., 2011 |
| Inferior & Superior Parietal Lobule | R | 7 | 3 | 34 | -54 | 46 | 18 (27) | 4728 | Cho et al., 2022; Aarts et al., 2010; Mizuno et al., 2008; Belayachi, et al. 2015; Bruening et al., 2018; Krebs et al., 2012; Orr et al., 2019; Vassena et al., 2014; Kostandyan et al., 2020; Boehler et al., 2014; Ivanov et al., 2012; Lallement et al., 2014; Leong et al., 2018; Paschke et al., 2015; Padmala et al., 2010; Padmala et al., 2011; Padmala et al., 2017; Wilbertz et al., 2014 |
| Inferior Frontal gyrus | L | 6 / 9 | 4 | -44 | 8 | 32 | 16 (20) | 3704 | Cho et al., 2022; Bahlman et al., 2015; Mizuno et al., 2008; Krebs et al., 2012; Soutschek et al., 2015; Vassena et al., 2014; Kostandyan et al., 2020; Kouneiher et al., 2009; Luethi et al., 2016; Chikara et al., 2018; Boehler et al., 2014; Leong et al., 2018; Paschke et al., 2015; Wilbertz et al., 2014; Padmala et al., 2010; Padmala et al., 2011 |
| | R | 13 | 5 | 34 | 20 | 2 | 11 (12) | 2080 | Cho et al., 2022; Westbrook et al., 2019; Mizuno et al., 2008; |

| | | | | | | | | | |
|---|---|---|---|---|---|---|---|---|---|
| Insula | | | | | | | | | Bruening et al., 2018; Krebs et al., 2012; Boehler et al., 2014; Ivanov et al., 2012; Lallement et al., 2014; Leong et al., 2018; Wilbertz et al., 2014; Padmala et al., 2011 |
| Insula | L | 13 | 6 | -32 | 20 | 6 | 10 (11) | 1656 | Cho et al., 2022; Westbrook et al., 2019; Mizuno et al., 2008; Belayachi, et al. 2015; Bruening et al., 2018; Krebs et al., 2012; Boehler et al., 2014; Lallement et al., 2014; Padmala et al., 2017; Wilbertz et al., 2014 |
| Middle Frontal Gyrus | L | 6 | 7 | -26 | 0 | 50 | 7 (8) | 1456 | Cho et al., 2022; Belayachi, et al. 2015; Vassena et al., 2014; Boehler et al., 2014; Taylor et al., 2004; Wilbertz et al., 2014; Padmala et al., 2010 |
| Inferior Frontal Gyrus | R | 9 | 8 | 46 | 10 | 26 | 7 (8) | 1400 | Cho et al., 2022; Orr et al., 2019; Kostandyan et al., 2020; Luethi et al., 2016; Chikara et al., 2018; Padmala et al., 2011; Padmala et al., 2017 |
| Thalamus | R | | 9 | 12 | -10 | 6 | 6 (7) | 1040 | Mizuno et al., 2008; Luethi et al., 2016; Boehler et al., 2014; Ivanov et al., 2012; Leong et al., 2018; Wilbertz et al., 2014 |

**Table S3.** The ALE meta-analysis contrasting cued versus un-cued demand levels. For each coordinate, region label, hemisphere (right, left or bilateral), Brodmann area, MNI coordinates, ALE maxima, *p* values, *Z* values, cluster size ($mm^3$), and number of studies are provided.

| **Brain Region** | **Hemi** | **Cluster No.** | **x** | **y** | **z** | **N Studies (n Foci)** | **Volume ($mm^3$)** | **Studies in Cluster** |
|---|---|---|---|---|---|---|---|---|
| *Cued > Un-cued* | | | | | | | | |
| Medial Frontal Gyrus, Superior Frontal Gyrus, Cingulate Gyrus | R/L | 1 | -4 | 16 | 46 | 9 (10) | 984 | Boehler etal, 2014; Lallement et al., 2014; Ivanov et al., 2012; Kostandyan et al., 2020; Orr et al., 2019; Padmala & Pessoa, 2011; Cho et al., 2022; Taylor et al., 2004; Westbrook et al., 2019 |
| *Un-cued > cued* | | | | | | | | |
| Inferior Frontal Gyrus, Precentral Gyrus, Middle Frontal Gyrus | R | 1 | 42 | 12 | 26 | 2 (2) | 912 | Chikara et al., 2018; Luethi et al., 2016 |
